# Protective functions of ZO-2/Tjp2 expressed in hepatocytes and cholangiocytes against liver injury and cholestasis

**DOI:** 10.1101/841080

**Authors:** Jianliang Xu, P. Jaya P. Kausalya, Noémi Van Hul, Matias J. Caldez, Shiyi Xu, Alicia Ghia Min Ong, Wan Lu Woo, Safiah Mohamed Ali, Philipp Kaldis, Walter Hunziker

## Abstract

**BACKGROUND & AIMS:** Tight junctions (TJs) establish tissue barriers that maintain osmotic homeostasis and, in the liver, isolate bile flow from the blood circulation. ZO-2/Tjp2 is a scaffold protein that tethers TJ transmembrane proteins to the actin cytoskeleton. Missense mutations in Tjp2 have recently been shown to cause progressive cholestatic liver disease in humans. However, the underlying mechanisms still remain elusive. To study the role of Tjp2 in cholestatic liver disease, we generated and characterized mice lacking Tjp2 in hepatocytes, cholangiocytes, or both.

**METHODS:** Tjp2 was inactivated in the mouse liver (both in hepatocytes and cholangiocytes) or hepatocytes or cholangiocytes only. Liver function tests were carried out by biochemical analysis of plasma and liver samples and liver tissue was evaluated by immunohistochemistry and histology. The mice were also subjected to cholic acid (CA) diet to assess their susceptibility to liver insults.

**RESULTS:** Deletion of Tjp2 in the mouse liver did not result in apparent changes in TJ structure and composition, but lead to progressive cholestasis with lower expression levels of the bile acid (BA) transporter ABCB11/Bsep and the detoxification enzyme Cyp2b10. Feeding a CA diet that is well tolerated by control mice caused severe cholestasis and necrotic liver injury in mice lacking hepatic Tjp2. Administration of a CAR agonist, TCPOBOP, protected these mice from CA induced injury by enhancing the expression of the detoxifying enzyme Cyp2b10 in hepatocytes. Mice lacking Tjp2 in only hepatocytes or in only cholangiocytes showed less severe CA diet induced liver injury.

**CONCLUSION:** Loss of Tjp2 from hepatocytes and cholangiocytes both contribute to progressive cholestatic liver disease and higher susceptibility to liver injury. In hepatocytes, Tjp2 exerts a protective role by regulating expression levels of BA transporters and detoxification enzymes. The mice may provide a new animal model for cholestatic liver disease linked to Tjp2 mutations in humans.

## INTRODUCTION

In mammals, the liver is a core metabolic hub responsible for major physiological functions including amino acid, carbohydrate and lipid metabolism, detoxification, and bile secretion. These processes require the coordinated function of several specialized cell types found in the liver, including hepatocytes and bile duct epithelial cells or cholangiocytes. Hepatocytes and cholangiocytes rely on TJs to establish tissue barriers that, for example, maintain osmotic homeostasis, or segregate bile from the blood circulation to form the blood-bile-barrier. TJs are furthermore critical in maintaining the polarized distribution of specific transporters and detoxification and metabolic enzymes to the apical and basolateral plasma membranes of hepatocytes and cholangiocytes. This spatial segregation contributes to the blood-bile barrier as it is critical for processes such as the directional collection of BA from the blood and their release into the bile by hepatocytes, or the concentration of bile in the bile ducts. TJ-dependent apico-basal polarity furthermore enables the selective protein and lipid compositions of the canaliculus and lumen facing cell surfaces of hepatocytes and bile duct epithelial cells, respectively, that provides resistance of these cells to the detergent effects of bile.

Cholestasis, a process whereby bile acids, bilirubin and liver enzymes, in particular AP, spill into the blood circulation, is a hallmark of many liver diseases ^1^. Cholestasis results from obstruction of bile flow or as a consequence of acute and chronic liver injury, which likely compromise the blood-bile barrier. Not surprisingly, alterations in TJs and/or blood-bile barrier function have been associated with a variety of cholestatic diseases and cholangiopathies. Further evidence for the critical role of TJs was the identification of genetic mutations inactivating the Tjp2 gene in cholestatic liver disease patients ^2-4^. Interestingly, mutations in Tjp2 were also found in Amish patients suffering from familial hypercholanaemia, a rare disease presenting with pruritus and elevated serum bile acids without liver disease. A symptomatic patient lacking the variant as well as several asymptomatic individuals homozygous for the variant were found, suggesting reduced penetrance and oligogenic inheritance.

The Tjp2 gene encodes for ZO-2/Tjp2, a cytosolic protein that has been extensively characterized as a structural component of TJs. Tjp2 tethers TJ integral membrane proteins, such as members of the Claudin family, to the actin cytoskeleton. In addition to this scaffolding role, Tjp2 has been implicated in different signalling pathways ^5^ and in sub-confluent or wounded cell monolayers redistributes from junctions to the nucleus ^6^.

In the present study we inactivated Tjp2 in the liver of mice and show the progressive development of cholestasis. These mice are also sensitized to liver injury by external insults, such as CA diet. Inactivating Tjp2 did not result in a gross alteration of the blood-bile barrier. Interestingly, however, changes in transcript levels of bile acid transporters and detoxification enzymes lead to increased liver bile acid levels which may not be adequately detoxified, thus inducing liver injury. While Tjp2 in both hepatocytes and cholangiocytes is important for liver function, inactivating Tjp2 in one or the other cell type showed differential effects on liver pathophysiology and susceptibility to injury.

## MATERIALS AND METHODS

### Mouse strains, genotyping

Animal experimentation was approved by the relevant IACUC and carried out in accordance to IACUC Protocol #171211. To delete Tjp2 in mouse hepatocytes, cholangiocytes or both, C57BL/6Tac Tjp2^F/F^ mice ^7^ were crossed with C57BL/6 Alb-Cre^ERT2^, Sox9-Cre^ERT2^ or Alb-Cre mice (Jackson Laboratory), respectively, and backcrossed into the Tjp2^F/F^ background. Genotyping was done on genomic DNA isolated from tail clippings amplified using primer-1 (5’-GTT CCT ATC CTG TTA GTT GGT AGT CC-3’) and primer-2 (5’-AAA GGG TCT CAT GTA GGT CAA GC-3’), yielding a 265 bp fragment for the wild-type allele and a 422 bp fragment for the conditional mutant allele. The absence of Tjp2/ZO-2 protein was verified by immunostaining and Western blot. C57BL/6Tac Tjp2^F/F^ littermate mice were used as controls and they showed no significant differences to a mixed cohort of C57BL/6Tac wild-type, C57BL/6Tac Alb-Cre, or C57BL/6Tac Alb-Cre Tjp2^F/+^ animals in terms of blood biochemistry at P60 and P300, or in susceptibility to 0.5% CA feeding (Supplemental Data 1). Tamoxifen (Sigma-Aldrich, St. Louis, MO) was dissolved in sunflower oil (Sigma-Aldrich, St. Louis, MO) to a concentration of 20 mg/ml and injected intraperitoneally at a dosage of 100 mg/kg for 5 days. No significant differences were detected between tamoxifen treated C57BL/6Tac Tjp2^F/F^ and C57BL/6Tac Alb-Cre^ERT2^ Tjp2^F/+^ or C57BL/6Tac Sox9-Cre^ERT2^ Tjp2^F/+^ animals in terms of liver to body weight ratio and blood biochemistry (data not shown).

### Cholic acid diet and TCPOBOP treatment

Were indicated, mice were fed with chow diet supplemented with 0.5% or 1% cholic acid (CA; Catalog no. TD.06026 and TD.10056; Harlan Teklad) for 7 days. Mice were fasted for 4 hours before sacrifice. To rescue the cholestatic liver injury, mice were injected intraperitoneally with either 3 mg/kg TCPOBOP (Sigma-Aldrich, St. Louis, MO) or vehicle (sunflower oil) daily for 5 days concomitant with the beginning of the cholic acid diet feeding.

### Histology

Freshly dissected liver was fixed in 4% paraformaldehyde overnight, processed, and embedded in paraffin. The paraffin block was sectioned to 5 μm sections and stained with hematoxylin and eosin, or Sirius red. The slides were viewed with a Zeiss Axio microscope, images were taken with a Zeisscam camera and representative images are shown.

### LacZ staining

Freshly dissected liver was frozen in OCT (optimal cutting temperature) compound and 10 µm thick sections were cut and mounted on slides. The slides were fixed in formalin for 10 minutes and then incubated in X-gal working solution at 37°C for 24 hours. After briefly rinsing in distilled water, the slides were counterstained with nuclear faster red for 3-5 minutes. Representative images are shown.

### Immunostaining

Paraffin sections were dewaxed and the antigen was retrieved by steaming the slides for 20 minutes in a 2100 Retriever (Pick Cell Laboratories). Sections were then stained with primary antibodies against ZO-1 (rat; Cat #R26.4C DSHB), ZO-2 (Tjp2) (rabbit; Cat #71-1400 Invitrogen), Claudin-1 (rabbit; Cat #71-7800 Invitrogen), Claudin-2 (rabbit; Cat #32-5600 Invitrogen), Claudin-3 (rabbit; Cat #34-1700 Invitrogen), Occludin (rabbit; Cat #71-1500 Invitrogen), VE-cadherin (Rat; Cat #550548 Pharmingen) or CK19 (rat; Cat #Troma III DSHB). Images were obtained using Zesis LSM800 confocal microscope and representative images are shown.

### Western Blot Analysis

Freshly dissected liver was homogenized with a mortar and lysed for 15 min in lysis buffer (50 mM Tris-HCl, pH7.5, 100 mM NaCl, 1 mM MgCl_2_, and 0.5% Triton X-100, supplemented with protease inhibitor cocktail and one PhosSTOP tablet per 10ml (Cat.# 04 906 837 01, Roche) on ice. Lysates were sonicated and cleared by centrifugation (13,000 × g for 15 min) at 4°C. Equal amounts of protein of the supernatant were fractionated by SDS-polyacrylamide gel electrophoresis and subjected to Western blot analysis using antibodies against ZO-1 (rabbit; Cat #61-7300, Invitrogen), ZO-2 (Tjp2) (rabbit; Cat #38-1100, Invitrogen), Claudin-1 (rabbit; Cat #ab15098, Abcam), Claudin-3 (rabbit; Cat #34-1700, Invitrogen), Occludin (rabbit; Cat #71-1500, Invitrogen), or GAPDH (Mouse; Cat #sc-47724, Santa Cruz). Images of representative blots are shown.

### Measurement of bile flow rate

In fasted mice under anesthesia the common extrahepatic bile duct was ligated and the gall bladder was cannulated. The animals were placed in a temperature-controlled incubator and bile secretions were collected over a one-hour period. Bile flow rate was assessed gravimetrically, assuming a bile density of 1 g/ml.

### Serum and tissue biochemical analysis

Serum alanine aminotransferase (ALT), aspartate aminotransferase (AST), alkaline phosphatase (AP), and bilirubin levels were analyzed using kits from Teco Diagnostics according to the manufacturer’s instructions. Plasma total bile acid levels were determined by Diazyme TBA Assay kit (Diazyme Laboratories). For liver bile acid levels, 100 mg liver tissue was ground in liquid nitrogen and mixed with 1 ml water. The mixture was sonicated, centrifuged and the supernatant collected for bile acid level determination using the Diazyme TBA Assay kit.

### mRNA extraction and Real-Time PCR

Samples from at least 3 independent mice for each group were isolated. For real-time PCR analysis, total mRNA was extracted from whole liver using RNeasy Mini Kit (250) from QIAGEN. Reverse transcriptions were performed with SensiFAST™ cDNA Sythesis Kit from Bioline. PCR reactions were carried out with the SensiFAST™, SYBR® No-ROX Kit (Bioline) in an Applied Biosysems cycler (Thermo Fisher Scientific). The respective forward and reverse primers for each RT-PCR are listed in Table 1.

**Table 1.**
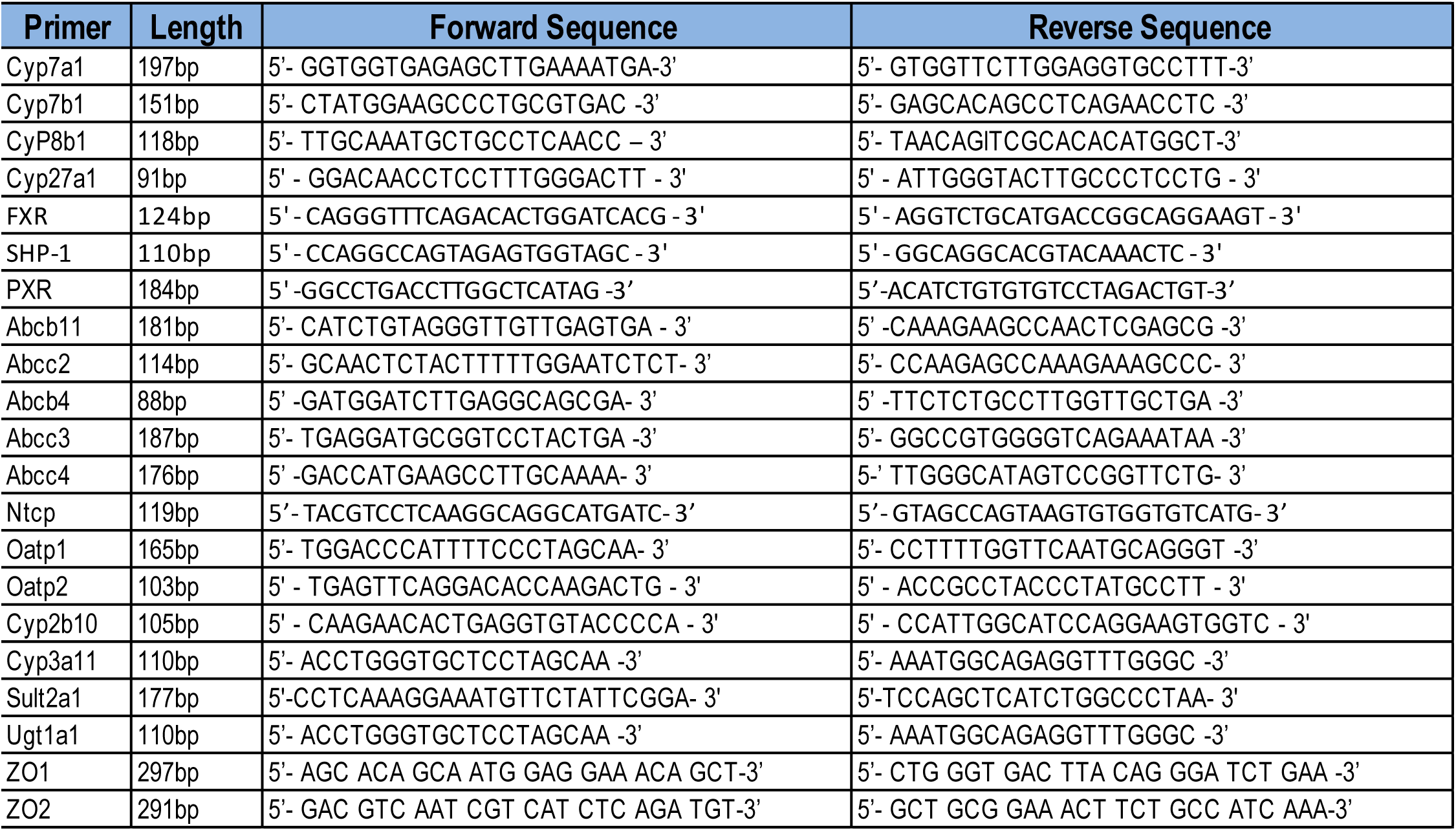

### Electron microscopy

Mouse livers were perfused with 2.5% glutaraldehyde in PBS via the inferior vena cava. After perfusion, the liver samples were cut into small pieces and kept in 2.5% glutaraldehyde for another 24 hours. After a rinse with 0.1 M PBS, the samples were postfixed in 1% osmium tetroxide for 1 h, rinsed in phosphate buffer, dehydrated in ethanol, and embedded in resin. Ultrathin sections were cut with a diamond knife, stained with uranyl acetate and lead citrate, and then viewed with a transmission electron microscope (JEM-1010). Representative images are shown.

### Statistical analysis

Samples from at least three independent experiments we used for real time qRT-PCR. For all other experiments, data from at least five independent animals or experiments were analyzed. The data are expressed as the mean ± SD. Paired Student’s t-test using Prism software (GraphPad) was used for statistical analysis and p<0.05 was considered significant.

## RESULTS

### Conditional deletion of Tjp2 in mouse liver hepatocytes and cholangiocytes

In order to inactivate Tjp2 in the mouse liver, Tjp2^F/F^ mice ^7^ were crossed to Albumin-Cre (Alb-Cre) mice to generate Tjp2^F/F^Alb-Cre animals, henceforth referred to as Tjp2 cKO mice, as well as control strains (Tjp2^F/F^, Tjp2^F/+^ Alb-Cre, and Alb-Cre). The albumin promoter is activated during liver development in common precursors that later give rise to both cholangiocytes and hepatocytes ^8^ and floxed genes in Alb-Cre mice are deleted in both hepatocytes and cholangiocytes ^9, 10^. As expected, Tjp2 cKO mice lacked Tjp2 in both of these cell types, but not in VE-Cadherin positive endothelial cells, as verified by immunofluorescence staining (Fig. 1A). The efficient deletion of Tjp2 in the liver was confirmed by Western blot analysis (Fig. 1B) and qRT-PCR (Fig. 1C). Neither localization nor expression levels for Tjp1 were significantly altered in the Tjp2 cKO liver at P60, with a slight reduction observed at P300 (Fig. 1C), suggesting that there was no compensatory increase of Tjp1 expression upon deletion of TJP2.

**Figure 1.**
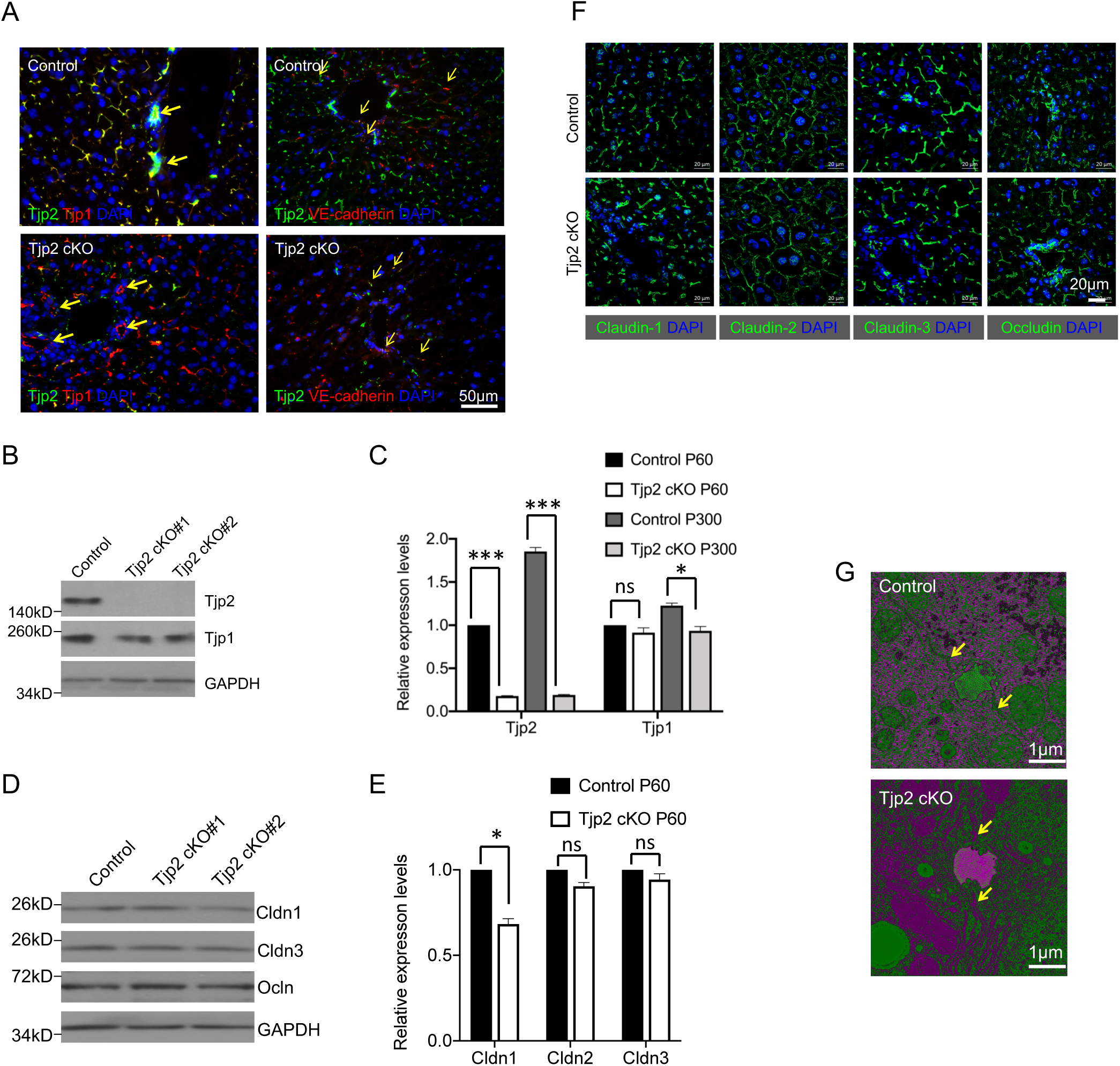
Conditional deletion of Tjp2 in the mouse liver and TJ structure. (**A**) Immunofluorescence microscopy of liver sections from control and Tjp2 cKO mice stained with antibodies against Tjp2, Tjp1 or VE-cadherin, showing absence of Tjp2 protein in both hepatocytes and bile duct cholangiocytes (thick arrows), but not in endothelial cells (thin arrows). Nuclei are labelled with DAPI. (**B**) Western blot analysis of whole liver lysates from a control and two independent Tjp2 cKO mice probed with antibodies to Tjp2, Tjp1 and GAPDH, confirming the absence of Tjp2 in the Tjp2 cKO liver, while Tjp1 protein levels are not affected. GAPDH was used as a control. (**C**) mRNA expression levels of Tjp2 and Tjp1 in whole liver lysates from P60 and P300 control and Tjp2 cKO mice determined by qRT-PCR. Residual Tjp2 mRNA levels likely reflect the presence of cell types other than hepatocytes and cholangiocytes, for example endothelial cells, which express Tjp2. (**D**) Western blot analysis of whole liver lysates from a control and two Tjp2 cKO mice at P60 probed with antibodies to Cldn1, Cldn3, and Ocln. GAPDH was used as a control. (**E**) mRNA expression levels of Cldn1, Cldn2 and Cldn3 in whole liver lysates from P60 control and Tjp2 cKO mice determined by qRT-PCR. Data in (D) and (E) are shown as mean ± SD, paired Student’s t-test p-value <0.05 was considered significant. *=p<0.05; **=p<0.005, ***=p<0.0005, ns=not significant (p>0.05) (**F**) Immunofluorescence microscopy of liver sections from P60 control and Tjp2 cKO mice stained with antibodies to Cldn1, Cldn2, Cldn3, and Ocln. (**G**) Electron microscopy images of liver sections from control and Tjp2 cKO mice showing the typical electron dense TJ structures in the vicinity of the bile canaliculi.

### Deletion of Tjp2 and expression and localization of key tight junction markers and structural appearance of the junctional complex

Tjp2 is a major structural component of the TJ and its absence may affect the structure, and as a consequence, the function of the TJs ^11-13^. While expression levels of Cldn1 were moderately lower, those of Cldn3 or Ocln were not significantly different in Tjp2 cKO as compared to control liver (Fig. 1D) when assessed by Western blot analysis or qRT-PCR. Frozen liver sections stained with antibodies Cldn1, Cldn2, Cldn3 and Ocln showed similar staining intensities and localization for these proteins in Tjp2 cKO and control samples (Fig. 1E). Deletion of TJP2 did not affect the structure of the bile canaliculi as delineated by markers such as Cldn1, Cldn2 and Cldn3. This was also evident by electron microscopy imaging, where the typical electron dense TJ plaques were observed proximal to the canaliculus between adjacent hepatocytes, both in control and Tjp2 deficient liver sections (Fig. 1F). Combined, this shows that, in absence of TJP2, TJP1 is sufficient to form normal bile canaliculi between hepatocytes. We conclude that hepatic deletion of Tjp2 does not alter the TJ structure and, together with the observed gradual progression of liver injury (see below), does not induce an acute dysfunction or breakdown of the hepatic TJ barrier.

### Tjp2 cKO mice are more prone to spontaneous liver injury and progressive liver fibrosis

Analysis of several physiological parameters indicated that mice lacking Tjp2 spontaneously developed mild liver injury over time. The liver to body weight ratio was significantly increased in 10 month old (P300) Tjp2 cKO mice (Fig. 2A). Liver biochemistry showed significantly elevated ALT and AST plasma levels in 2 month old (P60) Tjp2 cKO animals, and these were markedly higher at 10 months of age (Fig. 2B-C). H&E staining revealed more bile duct structures in 2 month old Tjp2 cKO mice as compared to controls, and a further increase with age (Fig. 2E), suggesting ductular reaction in response to liver injury. Despite variability between animals, Sirius Red staining showed an increase in liver fibrosis with aging in Tjp2 cKO livers (Fig. 2D and F).

**Figure 2.**
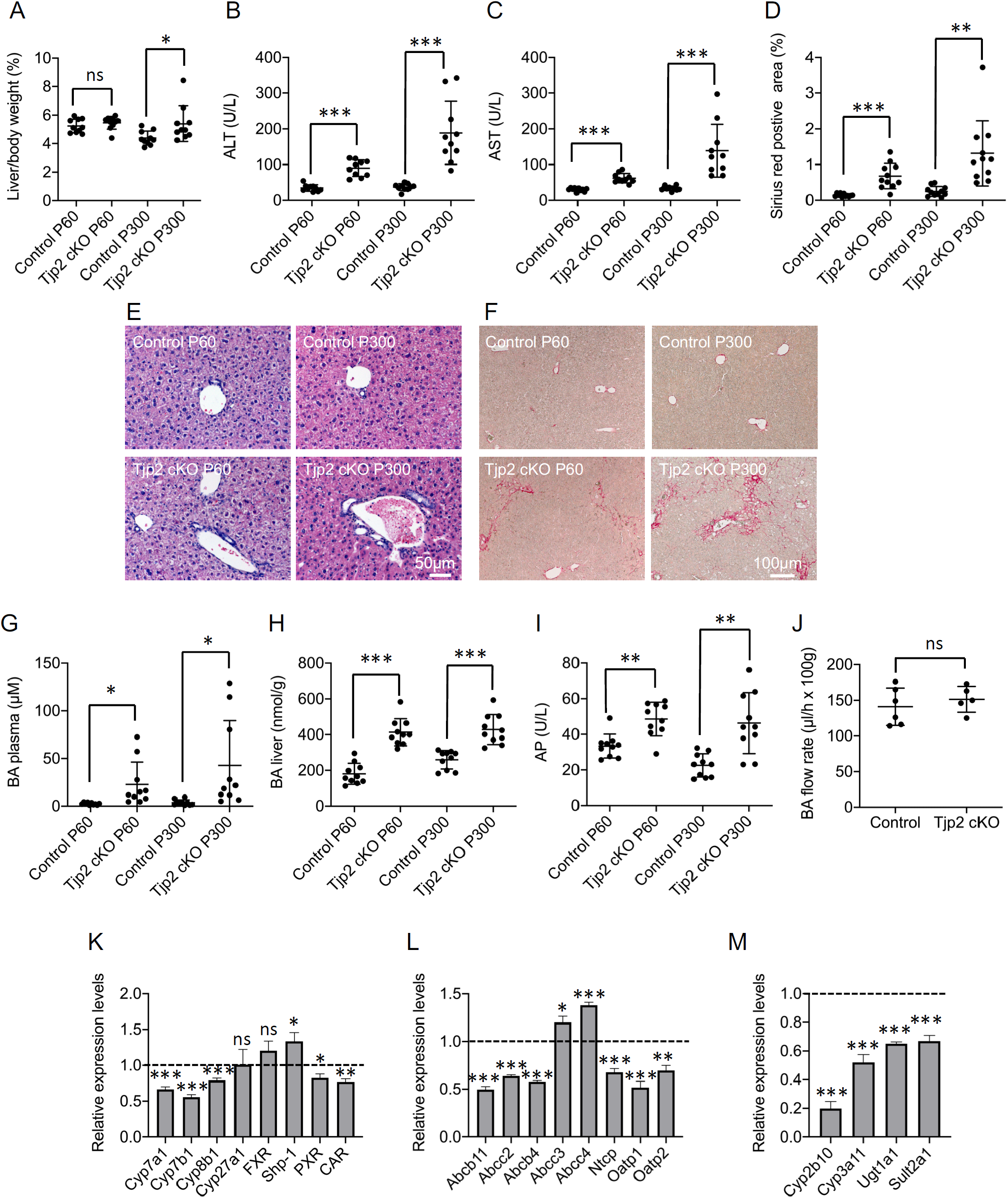
Blood and liver biochemistry and liver histology of control and Tjp2 cKO mice show progressive cholestasis in Tjp2 cKO mice. (**A**) Liver to body ratio. (**B, C**) ALT and AST plasma levels as markers for liver injury. (**D**) Quantification of Sirius red staining indicative of fibrosis from images as shown in panel F. H&E (**E**) and Sirius red (**F**) staining indicative of liver fibrosis. Plasma BA (**G**), liver BA (**H**) and AP levels (**I**). (**J**) Bile flow in livers of control and Tjp2 cKO mice. mRNA expression levels for BA synthesis genes (**K**), BA transporters (**L**) and detoxification enzymes (**M**) in the livers of control and Tjp2 cKO mice determined by qRT-PCR. Data are shown as mean ± SD, paired Student’s t-test p-value <0.05 was considered significant. *=p<0.05; **=p<0.005, ***=p<0.0005, ns=not significant (p>0.05).

### Conditional deletion of Tjp2 in mouse liver leads to increased bile acid levels and reduced expression of bile acid transporters and detoxification enzymes

In humans, mutations in Tjp2 have been associated with cholestasis ^3, 4^, which is characterized by increased BA levels in the circulation ^14^. Despite a large variability between individual animals, a tendency to higher BA levels in the plasma was observed for Tjp2 cKO mice, in particular at 10 months of age (Fig. 2G). These mice also presented with higher liver BA (Fig. 2H) and higher AP plasma (Fig. 2I) levels, consistent with cholestasis.

Cholestasis is the consequence of impaired bile flow due to obstruction of the biliary system, or deficiencies in bile acid uptake, conjugation, or excretion ^15^. To determine whether bile flow was altered in Tjp2 cKO mice, we cannulated the bile duct, collected the bile for one hour, and determined the excreted volume. The rate of bile flow in the Tjp2 cKO liver was comparable to that in controls (Fig. 2J), indicating that cholestasis in Tjp2 cKO mice did not result from impaired bile flow. Therefore, we next determined whether BA synthesis was increased by using qRT-PCR to monitor the expression levels of different enzymes involved in the conversion of cholesterol to BA (Fig. 2K). Cyp7a1, the rate limiting enzyme in the canonical pathway for bile acid synthesis ^16^, was repressed in the Tjp2 cKO liver, while mRNA levels for FXR and Shp-2, two suppressors of Cyp7a1, were higher. Cyp7b1 and Cyp8b1, two other enzymes in the BA synthesis pathway, were are also repressed. Although bile flow was not significantly affected in the liver of Tjp2 cKO mice, there were changes in the rate of bile acid synthesis. We next monitored the expression levels of BA transporters, which mediate the transfer of BA from hepatocytes into either the bile or the blood stream ^17^. Transcript levels of ABCB11/BSEP, the main transporter for BA into the bile canaliculus, were decreased in Tjp2 cKO livers as compared to controls (Fig. 2L). In contrast, mRNA expression levels of ABCC3 and ABCC4, both responsible for BA transport into the circulation, were elevated, whereas those of NTCP, Oatp1, and Oatp2, which remove BA from the blood stream, were decreased. Thus, these observations indicate that increased excretion of BA into the blood stream and decreased uptake could contribute to the observed cholestasis in Tjp2 cKO mice.

Cytotoxicity of BAs is diminished by modifications, such as hydroxylation, by detoxification enzymes ^18^. Exposure to ineffectively detoxified BA could contribute to chronic liver injury and cholestasis. Expression of two Phase I detoxification enzymes involved in BA hydroxylation, Cyp3a11 and Cyp2b10, was repressed in the absence of Tjp2 (Fig. 2M). Likewise, mRNA levels of the Phase II detoxification enzymes Sult2a1 and Ugt1a1 were reduced. This suggests that suboptimal detoxification of BA in the absence of Tjp2 could, over time, contribute to the observed liver injury.

In conclusion, the absence of Tjp2 in the mouse liver affects the expression of BA transporters and detoxification enzymes, likely promoting cholestasis and liver injury.

### Mice lacking Tjp2 are more susceptible to external hepatotoxic insults

Feeding a diet supplemented with cholic acid (CA), one of the two major bile acids produced by the liver, has been extensively used as a method to induce liver injury and cholestasis in mice ^19^. To test if the absence of Tjp2 from liver rendered mice more susceptible to CA diet, control and Tjp2 cKO mice were fed for 7 days a diet containing subacute levels of CA (0.5%), which is well tolerated by wild-type mice ^19^. Liver biochemistry and histopathology were assessed one week later. Compared to controls, the serum of Tjp2 cKO animals showed a yellow coloration (Fig. 3A), with significantly higher serum bilirubin (Fig. 3B) and AP (Fig. 3C), as well as serum and liver bile acid levels (Fig. 3D and E), consistent with cholestasis. H&E staining showed a normal liver histology in control livers, whereas Tjp2 cKO animals showed necrotic hepatic lesions (Fig. 3F-G) and elevated AST levels (Fig. 3H), indicative of liver injury. Indeed, qRT-PCR analysis revealed an increase in mRNA levels of Phase I and Phase II detoxification enzymes in response to 0.5% CA diet, which was abrogated in Tjp2 cKO animals (Fig. 3I). Expression of genes involved in bile acid synthesis and uptake was inhibited in both control and Tjp2 cKO liver (Fig. 3J). The CA diet-induced upregulation of the bile acid transporter ABCB11/BSEP in controls was not observed in Tjp2 cKO mice (Fig K). We conclude therefore that a failure to upregulate the expression of detoxification enzymes and ABCB11/BSEP contribute to the increased susceptibility to injury of the Tjp2 cKO liver.

**Figure 3.**
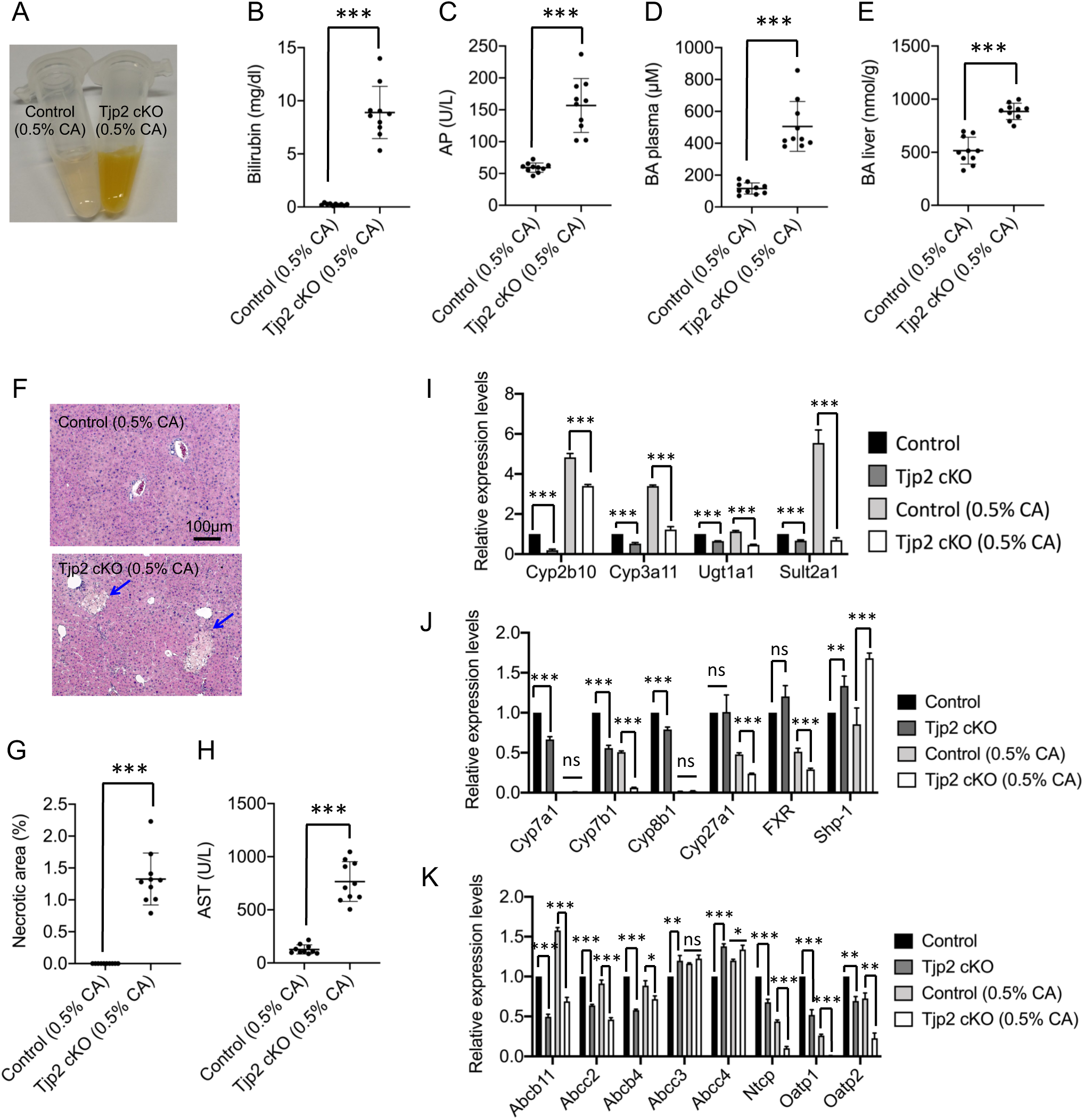
Tjp2 cKO mice are more susceptible to CA diet induced liver injury and cholestasis. (**A**) Plasma from control and Tjp2 cKO mice after a 7-day 0.5% CA diet. Bilirubin (**B**), AP (**C**), plasma BA (**D**), and liver BA (**E**) levels. (**F**) H&E staining showing liver necrosis (arrows) in CA diet fed Tjp2 cKO mice. (**G**) Quantification of necrotic area. (**H**) Plasma AST levels. mRNA expression levels for detoxification enzymes (**I**), BA synthesis genes (**J**), BA transporters (**K**) in the livers of control and Tjp2 cKO mice fed with normal or 0.5% CA diet as determined by qRT-PCR. Data are shown as mean ± SD, paired Student’s t-test p-value <0.05 was considered significant. *=p<0.05; **=p<0.005, ***=p<0.0005, ns=not significant (p>0.05).

### Stimulating detoxification enzyme expression in Tjp2 cKO mouse liver reduces liver injury

The transcriptional regulator CAR is a nuclear receptor that plays a central role in the sensing and clearance of toxins through the upregulation of genes encoding detoxification enzymes (e.g. Cyp2b10 and Cyp3a11) and bile acid transporters (e.g. ABCC3 and ABCC4) ^20^. We hypothesized that injection of the CAR agonist TCPOBOP ^21^ should therefore reduce the extent of the liver injury induced by 0.5% CA in Tjp2 cKO mice. As expected, TCPOBOP injection did not affect CAR expression levels but it lead to an increase in mRNA levels for several detoxification enzymes, with an over 100-fold increase of Cyp2b10 transcript levels (Fig. 4A). While expression levels of bile acid transporters secreting bile acids into the blood (e.g. ABCC3 and ABCC4) were also increased, ABCB11/BSEP, which secrete bile acids into the bile, were not affected. In agreement with the suppression of liver injury induced by 0.5% CA diet, H&E staining showed little if any necrosis in the liver of Tjp2 cKO mice treated with TCPOBOP when compared to vehicle-treated controls (Fig. 4B-C). Consistent with the enhanced expression of ABCC3 and ABCC4, plasma and liver bile acid levels were reduced (Fig 4D-E), as were those for AST (Fig. 4F) and bilirubin (Fig. 4G). TCPOBOP failed to improve the liver biochemistry values to a similar extent as observed in 0.5% CA fed control mice, suggesting that Tjp2 exerts additional, CAR-independent protective functions.

**Figure 4.**
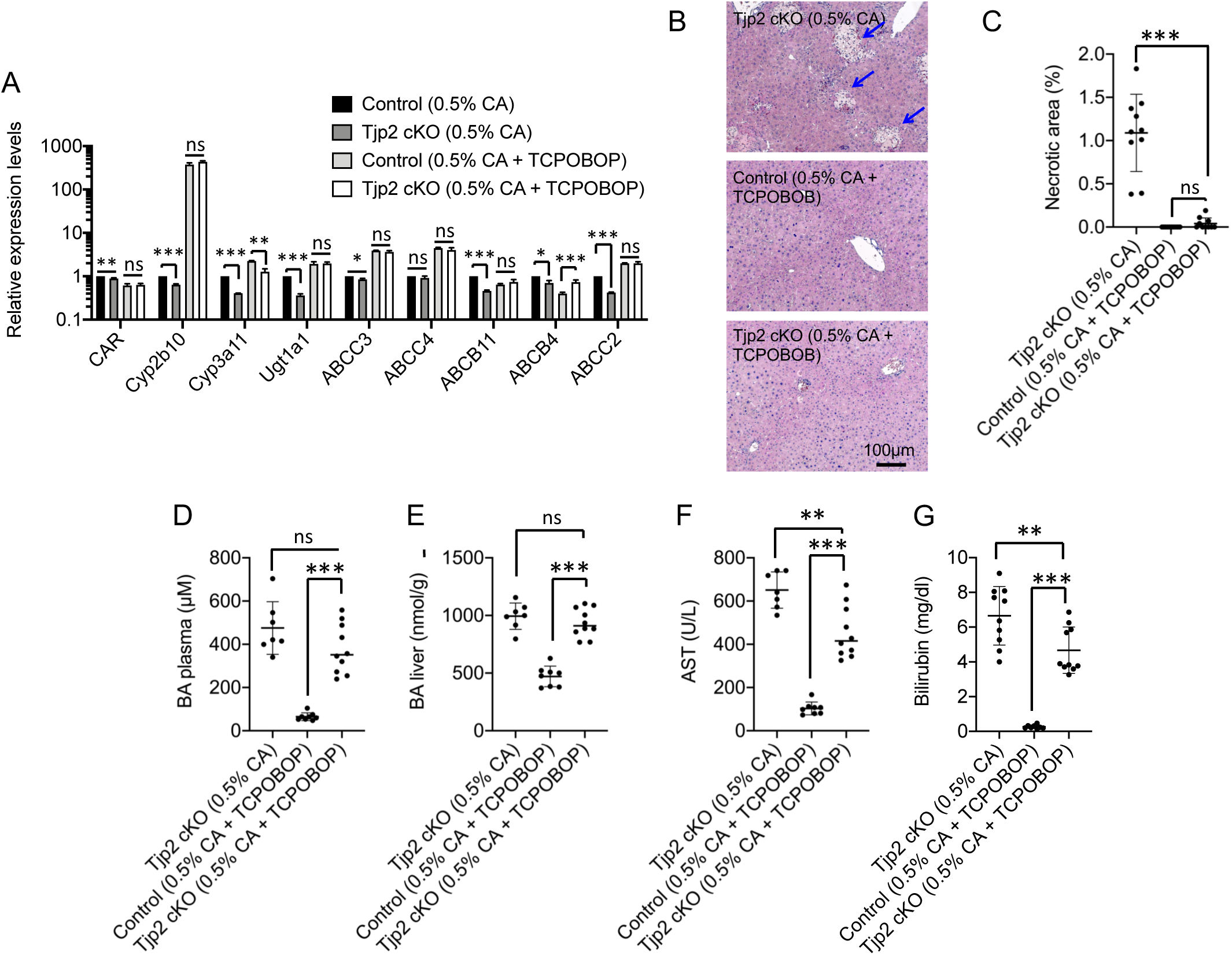
The CAR agonist TCPOBOP suppresses CA diet-induced necrotic injury in Tjp2 cKO mice.(**A**) mRNA expression levels for selected detoxification enzymes and BA transporters in control and Tjp2 cKO mice after a 5-day CA diet with or without daily TCPOBPOP injections, monitored by qRT-PCR. (**B, C**) TCPOBOP H&E staining and quantification of the necrotic area in livers of CA-diet fed control or Tjp2 cKO mice with or without TCPOBPOP administration. Plasma BA (**D**), liver BA (**E**), AST (**F**) and bilirubin (**G**) levels in CA-diet fed control or Tjp2 cKO mice with or without TCPOBOP treatment. Data are shown as mean ± SD, paired Student’s t-test p-value <0.05 was considered significant. *=p<0.05; **=p<0.005, ***=p<0.0005, ns=not significant (p>0.05).

### Conditional deletion of Tjp2 individually in hepatocytes or cholangiocytes

In the Tjp2 cKO mouse model used so far in this study, Tjp2 is inactivated in both hepatocytes and cholangiocytes (see above). To determine if the protective role of Tjp2 to liver injury requires Tjp2 expression in both of these cell types, or only in hepatocytes or cholangiocytes, we generated additional mouse models.

To inactivate Tjp2 in hepatocytes only, Tjp2^F/F^ mice were crossed with Albumin-Cre^ERT2 22^ mice to generate Tjp2^F/F^ Albumin-Cre^ERT2^ animals, referred to henceforth as Tjp2 icKO^HC^. Following five days of tamoxifen injection in 6-8 week-old mice, Cre expression was monitored by the activation of a lacZ reporter. While more than 99% of hepatocytes were LacZ positive, no LacZ activity was detected in bile duct structures in the vicinity of the portal vein, indicating efficient and hepatocyte-specific activation of Cre recombinase (Fig. 5A). This was corroborated by co-staining liver sections with antibodies to Tjp2 and the cholangiocyte marker CK19. While Tjp2 protein was no longer detected in CK19-negative hepatocytes following tamoxifen-mediated induction of Cre expression, Tjp2 was still present in CK19-positive cholangiocytes (Fig. 5B).

**Figure 5.**
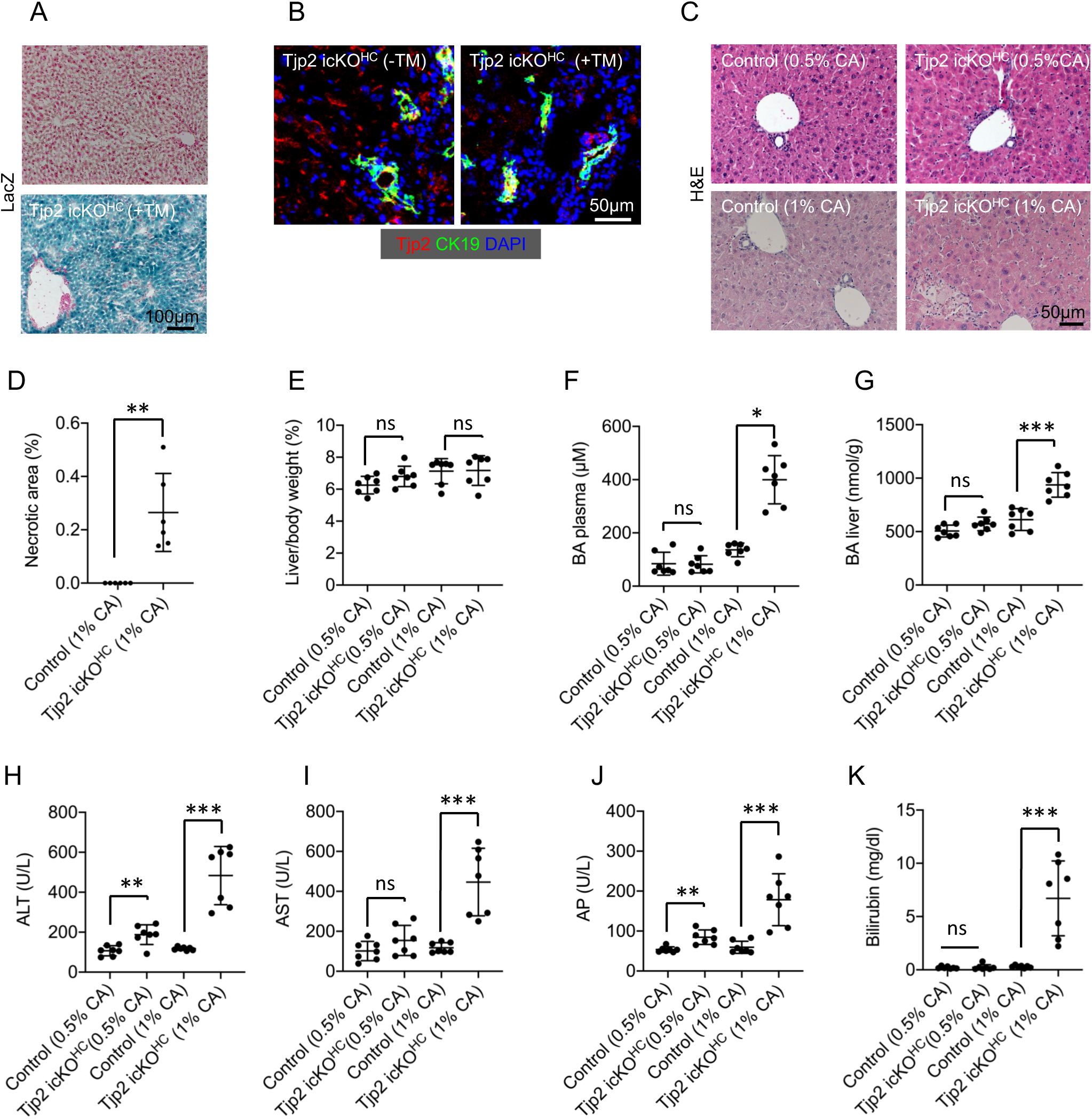
Mice with a hepatocyte-restricted inactivation of Tjp2 are less susceptible to CA-diet induced liver injury as compared to those lacking Tjp2 in both hepatocytes and cholangiocytes. (**A**) LacZ staining showing specific activation of the LacZ reporter gene in hepatocytes and not in cholangiocytes, indicative of a restricted deletion of Tjp2 in hepatocytes of Tjp2 icKO^HC^ mice. (**B**) Immunofluorescence microscopy of liver sections from Tjp2 icKO^HC^ mice treated or not with tamoxifen and stained with antibodies against Tjp2 and CK19, confirming the absence of Tjp2 protein in hepatocytes but not in CK19 positive cholangiocytes. Nuclei are labelled with DAPI. (**C**) H&E staining and quantification of necrotic area (**D**) showing reduced susceptibility of Tjp2 icKO^HC^ mice to CA-diet induced injury. (**E**). Liver to body weight ratios of control and Tjp2 icKO^HC^ mice fed 0.5% or 1% CA diet. Plasma (**F**) and liver (**G**) bile acid, ALT (**H**), AST (**I**), AP (**J**) and bilirubin (**K**) levels showing enhanced tolerance of Tjp2 icKO^HC^ animals to CA-diet induced injury (compare to respective parameters in Fig. 3 for Tjp2 cKO mice). Data are shown as mean ± SD, paired Student’s t-test p-value <0.05 was considered significant. *=p<0.05; **=p<0.005, ***=p<0.0005, ns=not significant (p>0.05).

To selectively inactivate Tjp2 in cholangiocytes, Tjp2^F/F^ mice were crossed with Sox9-Cre^ERT2 23^ mice to generate Tjp2^F/F^ Sox9-Cre^ERT2^ animals, referred to as Tjp2 icKO^CC^. In these mice, tamoxifen-induced, Cre-mediated activation of LacZ expression only occurred in bile duct cells in the vicinity of the portal vein and not in hepatocytes (Fig. 6A). As confirmed by immunofluorescence staining, Tjp2 protein in the Tjp2 icKO^CC^ liver was absent from CK19 positive cholangiocytes, but still expressed in the canalicular membrane of the hepatocytes (Fig. 6B).

**Figure 6.**
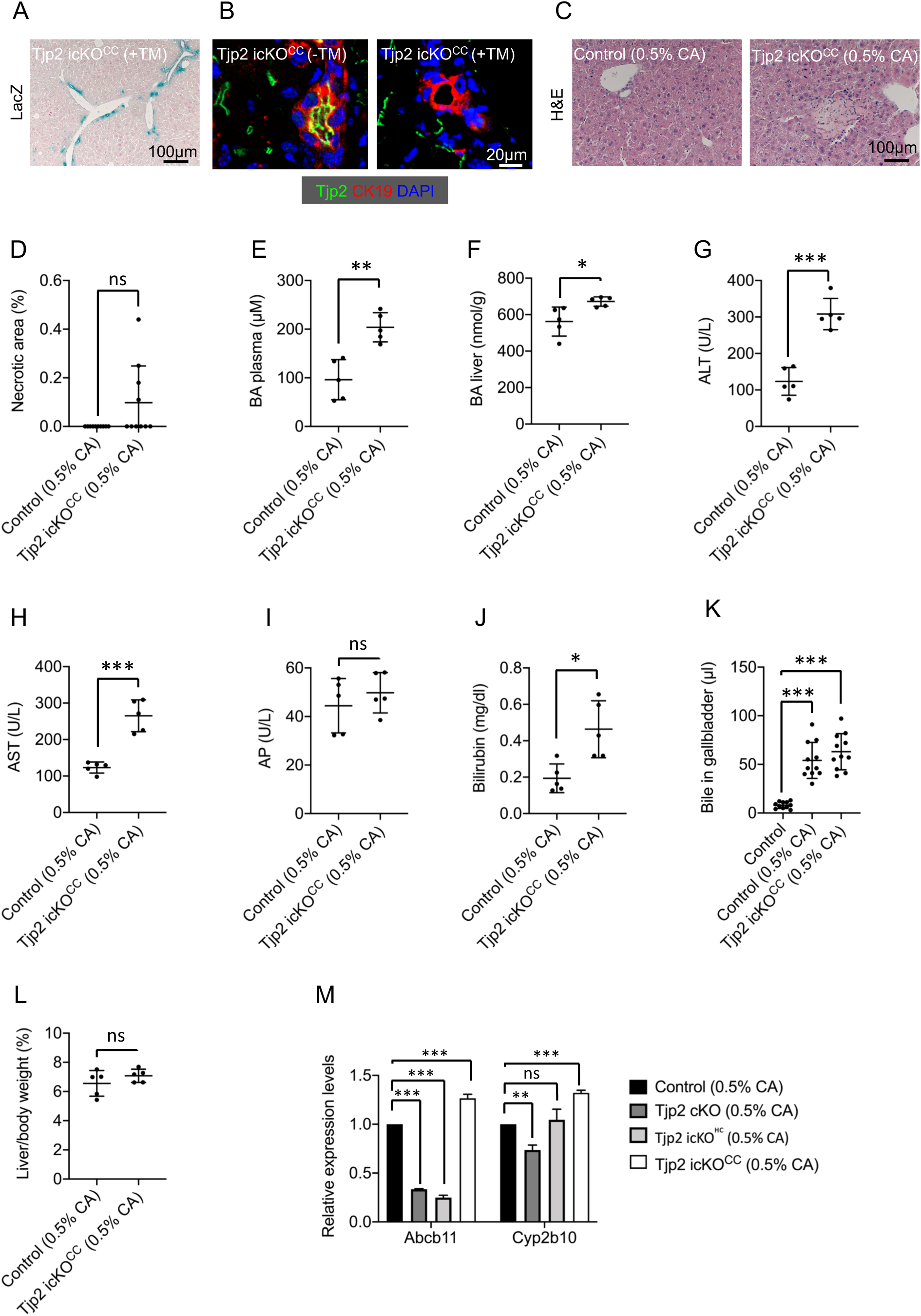
Deletion of Tjp2 in cholangiocytes contributes to the susceptibility to CA-diet induced liver injury. (**A**) LacZ staining showing specific activation of the LacZ reporter gene in bile ducts and not in hepatocytes, indicative of a restricted deletion of Tjp2 in cholangiocytes of Tjp2 icKO^CC^ mice. (**B**) Immunofluorescence microscopy of liver sections from Tjp2 icKO^CC^ mice treated or not with tamoxifen and stained with antibodies to Tjp2 and CK19, confirming the absence of Tjp2 protein in CK19 positive cholangiocytes but not in hepatocytes. Nuclei are labelled with DAPI. (**C**) H&E staining and quantification of necrotic area (**D**) of control and Tjp2 icKO^CC^ mice fed 0.5% CA diet. Liver (**E**) and plasma (**F**) BA, ALT (**G**), AST (**H**), AP (**I**) and bilirubin (**J**), and gall bladder BA (**K**) levels in control and Tjp2 icKO^CC^ mice fed 0.5% CA-diet. (**L**) Liver to body weight ratios. (**M**) mRNA expression levels for Abcb11 and Cyp2b10 in control, Tjp2 cKO, Tjp2 icKO^HC^ and Tjp2 icKO^CC^ mice fed with 0.5% CA diet monitored by qRT-PCR. Note the specific reduction of Abcb11 and Cyp2b10 expression levels only after deletion of Tjp2 in hepatocytes (e.g. in Tjp2 cKO and Tjp2 icKO^HC^ livers). Data are shown as mean ± SD, paired Student’s t-test p-value <0.05 was considered significant. *=p<0.05; **=p<0.005, ***=p<0.0005, ns=not significant (p>0.05).

### The loss of Tjp2 from cholangiocytes or hepatocytes both contribute to the increased sensitivity to liver injury

To determine whether the increased sensitivity of Tjp2 cKO mice to liver injury was due to the loss of Tjp2 in hepatocytes, cholangiocytes or both, we fed Tjp2 icKO^HC^ and Tjp2 icKO^CC^ mice 0.5% CA for one week, followed by 5 daily tamoxifen injections. While liver to body weight ratio was slightly increased (Fig. 5E), surprisingly little if any liver necrosis was apparent in Tjp2 icKO^HC^ mice (Fig. 5C). Increasing the CA concentration in the diet to 1% induced mild necrosis (Fig. 5C), with 0.2% of the tissue showing necrotic damage (Fig. 5D) as compared to over 1% in Tjp2 cKO mice fed with 0.5% CA (Fig. 3G). Plasma and liver bile acid levels (Fig. 5F-G), ALT (Fig. 5H), AST (Fig. 5I), AP (Fig. 5J) and bilirubin (Fig. 5K) levels were also similar to controls.

In contrast, feeding 0.5% CA diet to Tjp2 icKO^CC^ mice induced bile infarcts in ∼50% of animals (Fig. 6C). Average necrotic area (Fig. 6D), as well as plasma and liver bile acids levels (Fig. 6E-F) and ALT (Fig. 6G), AST (Fig. 6H) and AP (Fig. 6I) levels were significantly higher as compared to control mice, but lower than in Tjp2 cKO animals fed with 0.5% CA diet (compare to Fig. 3). Similarly, plasma bilirubin was significantly higher in 0.5% CA fed Tjp2 icKO^CC^ mice when compared to controls (Fig. 6J), but lower than in Tjp2 cKO animals (Fig. 3B). In addition, the volume of bile in the gallbladder was increased (Fig. 6K). Liver to body weight ratio was also higher in the Tjp2 icKO^CC^ mouse (Fig. 6L), probably reflecting induction of cell proliferation in response to liver injury. Similar to Tjp2 cKO, ABCB11/BSEP mRNA expression was strongly reduced in the Tjp2 icKO^HC^ liver, whereas Cyp2b10 levels were less affected (Fig. 6M). Interestingly, both ABCB11/BSEP and Cyp2b10 mRNA levels were increased in the Tjp2 icKO^CC^ liver. From these data we conclude that inactivation of Tjp2 exerts specific effects on hepatocytes and bile duct epithelial cells, as well as the physiological crosstalk between the two liver compartments.

## DISCUSSION

In humans, genetic mutations in Tjp2 are linked to cholestatic liver disease ^2-4^ and, in Amish, familial hypercholanaemia, a rare disease presenting with elevated serum bile acids without liver disease ^24^. Although the underlying mechanism has not been established, defects in the blood-bile barrier have been implicated since Tjp2 is a component of TJs. However, variations in the disease presentation and severity suggest that penetrance and possibly oligogenic inheritance may contribute. Here, we used mouse models to further characterize the role of Tjp2 in liver physiology and cholestasis. Due to embryonic lethality of the germline Tjp2 knockout mouse ^13^, we generated liver specific (Tjp2^F/F^ Albumin-Cre), hepatocyte-specific (Tjp2^F/F^ Albumin-CreERT2), and cholangiocyte specific knockout mice (Tjp2^F/F^ Sox9-CreERT2).

Tjp2 cKO mice lack Tjp2 in both hepatocytes and cholangiocytes, which both arise from albumin-expressing hepatoblast progenitors during liver development ^8^. The mice with hepatic Tjp2 deficiency developed mild progressive cholestasis. Expression levels and localization of Cldn1, Cldn2, Cldn3 or Ocln, major components of TJs, were moderately affected or indistinguishable from controls. Furthermore, transmission electron microscopy imaging showed that the typical electron dense TJ plaques in the vicinity of the canaliculus and between adjacent hepatocytes were maintained in the absence of Tjp2. Further arguing against an acute defect in the blood-bile barrier due to the absence of Tjp2 during liver development, liver biochemistry parameters were only mildly altered in 2 month old animals and had only progressed moderately by 10 months of age. Furthermore, while Tjp2 cKO mice fed with 0.5% CA diet developed severe cholestasis, Tjp2 icKO^HC^ animals fed this diet showed a comparably mild phenotype, similar to that observed in 0.5% CA fed controls. Thus, although Tjp2 is absent from the TJs of the bile canaliculi of both Tjp2 cKO and Tjp2 icKO^HC^ mice, the CA diet induced cholestasis differed greatly in severity and therefore cannot simply reflect a breakdown of the canalicular TJ barrier.

Impaired bile flow can increase the pressure in the biliary system and lead to obstructive cholestasis ^14^. However, bile flow rates where similar in the Tjp2 cKO mice as compared to controls. Synthesis, excretion and uptake of bile acids is tightly regulated and imbalances in these processes can lead to cholestasis ^25^. The mRNA for Cyp7a1, the rate limiting enzyme for bile acid synthesis in the mouse ^16^, was strongly repressed in the Tjp2 cKO mouse, probably due to negative feedback from the higher liver and plasma bile acid levels. Expression of other genes involved in bile acid synthesis was also inhibited, while their transcriptional repressors FXR and in particular Shp-1 were upregulated in the liver lacking Tjp2. Bile acids discharged into the small intestine are resorbed into the blood stream in the ileum and once they reach the liver are taken in by hepatocytes via bile acid transporters. The mRNA levels for several bile acid intake transporters, such as Ntcp, Oatp1 and Oatp2, were reduced in Tjp2 cKO mice and, while this might contribute to the higher plasma bile acid levels, lower intake from the circulation unlikely accounts for the observed cholestasis. Reduced excretion of bile acids from hepatocytes into the bile may, however, result in the observed higher liver bile acid levels in Tjp2 cKO mice. Indeed, mRNA levels for ABCB11/BSEP, the main bile acid transporter for the excretion of bile acids form hepatocytes into the bile, were reduced in the Tjp2 cKO mouse liver. When the mice were fed with 0.5% CA diet, the liver bile acid levels increased in control mice and even more so in Tjp2 cKO mice. CA diet lead to an increase in ABCB11/BSEP mRNA levels in control mice, but less so in Tjp2 cKO animals. ABCC3 and ABCC4 are responsible for transporting bile acids from hepatocytes into the blood stream. An increase in their expression in Tjp2 cKO mouse liver and might thus represent a rescue mechanism to reduce liver bile acid levels. Therefore, reduced ABCB11/BSEP expression might be a main contributor to cholestasis in Tjp2 cKO mice. In agreement with this interpretation, ABCB11/BSEP knockout mice fed with 0.5% CA diet develop severe cholestasis ^19^, similar to that observed in Tjp2 cKO mice. In addition, mRNA levels of detoxification enzymes, in particular Cyp2b10, which hydroxylates bile acids to increase hydrophilicity and reduce toxicity, were lower in Tjp2 cKO mice. The observed progressive development of fibrosis and eventually necrosis likely results from the chronic exposure of the liver to higher levels of bile acids that have not been optimally detoxified. In response to 0.5% CA diet, the liver of control animals showed an increase in detoxification enzyme levels, a response that was lacking in the Tjp2 cKO mice. Injection of these mice with the CAR-agonist TCPOBOP resulted in a greater than hundred fold increase in mRNA levels of the downstream gene Cyp2b10. This increase in the detoxification enzyme levels was sufficient to prevent liver necrosis in Tjp2 cKO mice fed with 0.5% CA diet, despite the fact that liver and plasma bile acid levels were not dramatically lowered by the agonist. Thus, liver injury in the Tjp2 cKO mouse may relate to the high lev levels of non-detoxified bile acids and enhancing the expression of detoxification enzymes, in particular Cyp2b10, is sufficient to inhibit this pathological process. Since CAR dependent gene transcription is enhanced by activation of Yap ^26^ and Tjp2 functionally associates with Yap/Taz ^27^, it may be possible that Tjp2 modulates the crosstalk between CAR and Yap.

Global inactivation of Tjp2 in the mouse results in embryonic lethality ^13^, suggesting developmental and/or tissue specific species differences for the importance of Tjp2. In contrast to the mild progressive cholestasis observed in Tjp2 cKO mice, a more severe progression of the disease has been reported for patients with genetic mutations that inactivate Tjp2. A reduction in Cldn1 protein levels in the liver of these patients was also observed, albeit to a lesser extent, in the Tjp2 cKO mice. There were no marked alterations in expression levels or localization of the other TJ components analyzed in the Tjp2 cKO liver. Changes in the expression of other TJ components or the ultrastructure of TJs, which have not been analyzed in in affected cholestatic patients, could account for the more severe phenotype compared to the Tjp2 cKO mice. It is also not know if expression of detoxifying enzymes or bile acid transporters, as seen in the mouse, are altered in these patients. Importantly, there are also significant differences in bile acid metabolism between human and mouse, which could explain differences in phenotypes between hepatic Tjp2 deficient mice and cholestatic patients with Tjp2 mutations. Enzyme activities that affect primary bile acid synthesis, conjugation and metabolism differ between human and mouse ^28-31^. In contrast to the mouse, most BA in human are FXR agonists ^32^. These differences result in a higher hydrophilicity and thus lower toxicity of the mouse bile compared to the human counterpart. Furthermore, behavioural differences such as coprophagy by rodents, or different metabolic modifications of BA by the human and mouse gut microbiome ^33, 34^, can influence BA profiles. Given the observed effects on BA transporters and detoxification enzymes in the mouse with hepatic Tjp2 deficiency, it will be of interest to determine if similar changes occur in cholestatic patients carrying Tjp2 inactivating mutations. If so, enhancing BA detoxification may provide a possible therapeutic approach for these patients.

The analysis of Tjp2 icKO^HC^ and Tjp2 icKO^CC^ mice revealed distinct contributions of the absence of Tjp2 in hepatocytes or cholangiocytes, respectively, to the pathophysiology observed in Tjp2 cKO mice, where Tjp2 was ablated in both cell types. While Tjp2 cKO mice developed severe cholestasis when fed with 0.5% CA diet, animals lacking Tjp2 only in hepatocytes or cholangiocytes tolerated this diet much better. Tjp2 icKO^HC^ mice started to develop minor liver necrosis when fed a 1% CA diet, while minor necrosis was already observed in Tjp2 icKO^CC^ animals provided with the 0.5% CA diet. Accordingly, liver bile acid levels after 0.5% CA diet in Tjp2 cKO mice were almost twice those in controls, but only moderately elevated in animals lacking Tjp2 in hepatocytes or cholangiocytes. Both. Tjp2 icKO^HC^ mice fed with 1% CA and Tjp2 icKO^CC^ fed with 0.5% CA showed necrotic liver lesions. However, plasma bilirubin levels were strongly elevated in Tjp2 icKO^HC^, but only moderately so in Tjp2 icKO^CC^ animals. One possibility is that in Tjp2 icKO^CC^ animals cholestasis and a larger gallbladder bile volume increase the pressure in the biliary system, leading to small ruptures of the bile ducts and leakage of bile into the interstitial space.

In conclusion, we show that Tjp2 plays a protective role in the liver. Liver specific inactivation of Tjp2 results in progressive cholestasis and liver injury. This likely does not reflect major effects on the TJs or the blood-bile barrier, but rather changes in the expression of genes for bile acid transporters and detoxification enzymes, in particular ABCB11/BSEP and Cyp2b10, respectively. Reduced excretion of bile acids from hepatocytes into the bile due to suppression of ABCB11/BSEP, combined with reduced levels of the detoxification enzyme Cyp2b10, may result in the accumulation of high levels of poorly detoxified bile acids in the liver, resulting in the observed pathology. How Tjp2 modulates the expression of ABCB11/BSEP and Cyp2b10, in particular in response to injury, remains to be elucidated.

## Abbreviations

Alb: albumin;
ALT: Serum alanine aminotransferase
AST: aspartate aminotransferase
AP: alkaline phosphatase
BA: bile acid
CA: cholic acid
CAR: constitutive androstane receptor
cKO: conditional knock-out,
Cre: Cre recombinase
FXR: farnesoid X receptor
icKO: inducible conditional knock-out
icKO^CC^: inducible cholangiocyte-specific knock-out
icKO^HC^: inducible hepatocyte-specific conditional knock-out
Ocln: occludin
TCPOBOP: 1,4-bis[2-(3,5-dichloropyridyloxy)]benzene
TJ: tight junction
ZO: zonula occludens
ZO-2: zonula occludens-2

## ACKNOWLEDGEMENTS

We thank the IMCB Advanced Molecular Pathology Lab for help in processing paraffin blocks, the Electron Microscopy Unit of the A*STAR Microscopy Platform for performing the electron microscopy and David Liebel for helpful discussions regarding liver sample preparation. This study was supported by the Biomedical Research Council of the Agency for Science, Technology and Research (A*STAR), Singapore, and grant OFIRG15nov120 from the National Medical Research Council, Singapore.

---

**Supplemental Figure 1.**
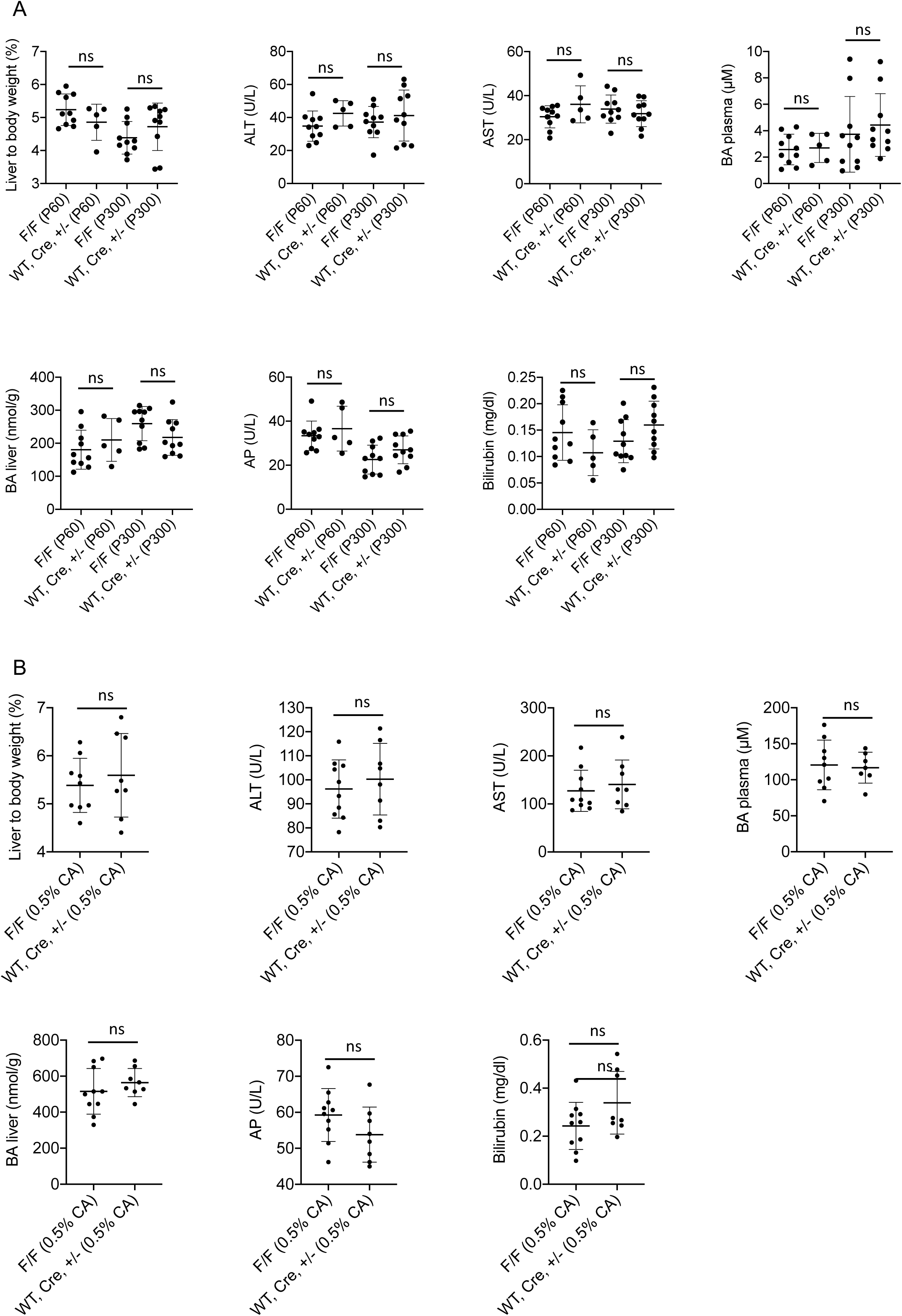
Blood and liver biochemistry for different cohorts of control mice. (**A)** Blood and liver biochemistry was carried out for tamoxifen-treated C57BL/6Tac wild-type, C57BL/6Tac Alb-Cre, C57BL/6Tac Alb-Cre Tjp2^F/+^ or C57BL/6Tac Tjp2^F/F^ animals at P60 and P300. The values for C57BL/6Tac wild-type, C57BL/6Tac Alb-Cre, or C57BL/6Tac Alb-Cre Tjp2^F/+^ were combined and compared to those of C57BL/6Tac Tjp2^F/F^ mice, used as controls in the study (e.g. Fig. 2A-C and G-I). (**B**) Blood and liver biochemistry were carried out for tamoxifen-treated C57BL/6Tac wild-type, C57BL/6Tac Alb-Cre, C57BL/6Tac Alb-Cre Tjp2^F/+^ or C57BL/6Tac Tjp2^F/F^ animals fed normal or CA-supplemented diet and compared to the C57BL/6Tac Tjp2^F/F^ mice used as controls (Fig. 3B-E and H). Data are shown as mean ± SD, paired Student’s t-test p-value <0.05 was considered significant. *=p<0.05; **=p<0.005, ***=p<0.0005, ns=not significant (p>0.05).

